# Mapping QTL associated with resistance to avian oncogenic Marek’s Disease Virus (MDV) reveals major candidate genes and variants

**DOI:** 10.1101/2020.07.22.215582

**Authors:** Jacqueline Smith, Ehud Lipkin, Morris Soller, Janet E. Fulton, David W. Burt

## Abstract

Marek’s Disease (MD) represents a significant global economic and animal welfare issue. Marek’s Disease Virus (MDV) is a highly contagious oncogenic and highly immune-suppressive alpha-herpes virus, which infects chickens, causing neurological effects and tumour formation. Though partially controlled by vaccination, MD continues to have a profound impact on animal health and on the poultry industry. Genetic selection provides an alternative and complementary method to vaccination. However, even after years of study, the genetic mechanisms underlying resistance to MDV remain poorly understood. The MHC is known to play a role in disease resistance, along with a handful of other. In this study, one of the largest to date, we used a multi-facetted approach to identify QTL regions (QTLR) influencing resistance to MDV, including an F_6_ population from a full-sib advanced intercross line (FSIL) between two elite commercial layer lines differing in resistance to MDV, RNA-seq information from virus challenged chicks, and genome wide association study (GWAS) from multiple commercial lines. Candidate genomic elements residing in the QTLR were further tested for association with offspring mortality in the face of MDV challenge in 8 pure lines of elite egg-layer birds. Thirty-eight QTLR were found on 19 chicken chromosomes. Candidate genes, miRNAs, lncRNAs and potentially functional mutations were identified in these regions. Association tests were carried out in 26 of the QTLR, using 8 pure lines of elite egg-layer birds. Numerous candidate genomic elements were strongly associated with MD resistance. Genomic regions significantly associated with resistance to MDV were mapped, and candidate genes identified. Various QTLR elements were shown to have strong genetic association with resistance. These results provide a large number of significant targets for mitigating the effects of MDV infection on both poultry health and the economy – whether by means of selective breeding, improved vaccine design or gene-editing technologies.

**Author summary:** Marek’s Disease has a huge impact on the global poultry industry in terms of both animal welfare and economic cost. For many years, researchers have sought to identify the genes underlying resistance to Marek’s Disease Virus (MDV). However, this is a complex trait with each genetic locus having a small effect, so identifying causal genes and variants is no easy task. To date, it is known that the MHC confers differing susceptibility/resistance. A few other non-MHC genes have also been implicated in disease resistance, although based on experimental inbred lines and not representing real world commercial poultry. Using an F_6_ intercross population of birds with differences in MDV survival, we have identified many regions of the genome involved in resistance and highlighted candidate genes, miRNAs and lncRNA. Access to DNA from phenotyped birds spanning 15 generations of 8 elite commercial lines has provided a unique opportunity for us to show genetic association of markers in these transcripts with MDV survival. This genetic study, the largest to date, provides novel targets for mitigation of Marek’s Disease within the poultry industry. This could be through selective breeding strategies, improved vaccine design or future gene editing technologies.

## Introduction

Marek’s Disease (MD) represents a significant global economic and animal welfare issue. This immunosuppressive disease is responsible for an estimated 2 billion USD annual economic loss to the global poultry industry [1], through bird mortality, lost egg production and vaccination costs. The virus responsible, Marek’s Disease Virus (MDV), is an alpha-herpes virus that initially infects B-cells, experiences a latency period and can then proceed to develop as an oncogenic disease after infection of T-cells [2]. Furthermore, if birds do not succumb to MDV itself, they are often left severely compromised with secondary infections such as *E. coli* [3]. The commercially utilized vaccines are not sterilizing vaccines. They prevent the formation of tumours, but do not prevent infection by MDV or shedding of the pathogenic virus [4]. Both vaccine and pathogenic MDVs are found in vaccinated flocks, resulting in the emergence of increasingly more virulent strains [5]. As more virulent strains emerge, vaccine treatments are becoming less and less effective [6].

In the race to keep ahead of this viral evolution, genetic selection is a possible tool to aid in breeding for viral resistance. Indeed, genetic selection following multiple generations of progeny challenge has been shown to improve survival in commercial chicken populations [7]. Selection in the poultry industry works relatively well, but knowing the causative elements for resistance would provide a route for much more precise selection. However, the underlying genetic changes, and the genes and specific variants impacting this genetic resistance are still largely unknown. For decades, researchers have sought to identify the genes responsible for MDV resistance, with limited success. It has become clear that many QTL/genes are involved in the resistance phenotype, with resistance not being a simple trait with one or a few major genes but a classic quantitative trait with many genes of small effect, thus making it difficult to identify causal variants [8].

Marek’s Disease is also of interest from a clinical perspective, as it can serve as a model for human lymphoma. MDV-induced lymphomas are found to over-express the Hodgkin’s disease antigen CD30, with expression correlating with the viral Meq oncogene [9]. Indeed, after infection, the chicken CD30 promoter has also been shown to be hypo-methylated [10]. Identifying the genetic mechanisms underlying MDV resistance is therefore not only of great importance to the poultry industry but will also have implications for increasing our understanding of human cancers.

Different MHC haplotypes are known to confer different susceptibilities to the virus [11–13]. Studies have also reported the influence of non-MHC regions on resistance to MDV [14]. Several studies have implicated specific genes in MDV resistance/susceptibility. These include *GH1* [15], *SCYC1* [16], *SCA2* [17], *IRG1* [18], *CD79B* [19] and *SMOC1* and *PTPN3* [20]. These studies are usually done, not with relevant commercial lines, but with experimental or inbred lines and examine whole tissues, although recent work has investigated the host response to MDV in specific cells such as macrophages, which are an early target for the virus [21,22]. Recent studies on the role of lncRNAs [23,24] and miRNAs (in both the host and virus) have also been carried out [25–27], including study of serum exosomes from lymphoma-bearing birds [28]. In addition, the role of epigenetics in resistance to MDV has been studied, with regions of differential methylation between susceptible and resistant lines of birds highlighted [29,30].

Here, the availability of large-scale, phenotyped commercial populations, genome wide analysis technologies and an F_6_ advanced intercross line [31], has given us the opportunity to carry out, for the first time, a high-resolution analysis of genes underlying MDV resistance in commercially relevant populations. We use multiple genetic resources at our disposal, including the F_6_ population of an advanced full-sib inter-cross line (FSIL) previously analysed in a low-resolution study for MD resistance using microsatellite markers to identify genomic regions associated with survival following MD challenge [31]. Genomic DNA of the original 10 founder individuals and the subsequently produced F_6_ was available for fine mapping through genome sequencing, and/or genotyping using a genome-wide 600K SNP chip [32]. Furthermore, an extensive multi-generation (15) and multi-line (8) collection of DNAs from progeny-challenged males was available to further examine candidate genes and related variants associated with survival in the face of MDV infection.

In this report we reveal for the first time MD as a true complex trait, controlled by many QTL. Integration of multiple lines of evidence (F_6_, multi-generation/multi-line collection, host gene expression responses to viral infection, genome annotations, etc.) on a large scale enabled a high-resolution analysis that predicted mutations within genes, miRNAs and lncRNAs highly associated with MDV response in commercial egg production lines. This analysis not only provides new markers for MD resistance but also reveals more about the biology behind the mechanism of MDV susceptibility, information that should lead to more precise selection strategies in the future.

## Materials and methods

### Experimental animals

#### Full Sib Advanced Inter-cross Line (FSIL)

F_6_ birds from the FSIL were used to map QTL affecting MD resistance. The development of the FSIL F_6_ challenge population has been previously described [31,33,34]. It was initiated from a cross of two partially inbred commercially utilized elite White Leghorn lines, known to differ in their resistance to MDV. Five independent FSIL families were developed and expanded over five generations. In all five families, the male parent was from the more resistant line and the female parent was from the less resistant line.

At the F_6_ generation, 1,615 chicks were challenged with vv+ MDV strain 686 following the protocol of Fulton et al. [7]. The experiment was carried out in two hatches; it was terminated at 152 days of age in the first hatch, and after 145 days of age in the second. Resistance was measured as survival time (age at death). For birds that survived to the end of the experiment, survival time was taken as age at end of the experiment. To standardise the two hatches, survival for all birds that survived to end of experiment was set to 149 days. This measure of resistance was used in all association tests.

This population was segregating for two MHC haplotypes (B^2^ and B^15^). Given the known strong association of MHC-type with MD resistance, all analyses were done within MHC-type within each family.

#### Pure line Pedigreed sire populations with daughter progeny records for MDV mortality

As part of the routine selection process within the Hy-line (HL) breeding program, individual males were mated to multiple females to produce 30 half-sib female progeny per sire. The dams were MD vaccinated following normal industry practices. Progeny were vaccinated at 1 day of age with HVT/SB1 and at 7 days of age inoculated subcutaneously with 500 PFU of vv+ MDV (provided by Avian Disease and Oncology Lab, East Lansing MI). Mortality was recorded from 3 to 17 weeks of age (termination of experiment), as described in Fulton et al. [7]. The sire MD tolerance phenotype is the proportion of mortality among the daughters upon MD challenge, with no signs of MD at the termination of the experiment.

Data were available from 15 generations of 8 elite lines (not the same generations in all lines), with 1,081 to 1,393 males per line having MD progeny mortality data, for a total of 9,391 males. The lines included three different egg production breeds, namely five White Leghorn lines, two White Plymouth Rock lines and one Rhode Island Red line. Pedigreed sires and their progeny from these pure lines were used to test functional genomic elements (genes, miRNAs, lncRNAs in the mapped QTL regions (QTLR)), for association with MD progeny mortality, Also tested were a group of singleton coding SNPs predicted to have deleterious effects on protein structure and function (henceforth, potentially functional mutations, see below for details).

### Genomic sequencing of founder birds of the FSIL

Genome sequence information was produced for the 10 F_0_ founder individuals of the F_6_ population. This was used to identify variants that differed in the genomes of the parental birds and may have functional impact. Sequencing the DNA from the 10 F_0_ founder birds was carried out by the Edinburgh Genomics sequencing facility (Edinburgh, UK). Samples were prepared for sequencing using 1 ug of genomic DNA following the TruSeq PCR free kit protocol (Illumina, FC-121-3001). Resulting libraries were quality checked on an Agilent DNA 1000 bioanalyzer (Agilent Technologies, South Queensferry, UK) and then clustered onto HiSeq Rapid V2 flow cell at a concentration of 15 pM. Sequencing was carried out on an Illumina Hiseq 2500 using the HiSeq Rapid v2 SBS reagents (Illumina, Little Chesterford, UK), for 150 cycle paired end reads. Each of the 10 samples was sequenced to around 15x coverage. Quality of sequences was determined using FastQC [35], and mapping to the chicken reference genome (Galgal4) was carried out using BWA (v0.7.0) [36]. The resulting bam file was sorted with Samtools (v0.1.18) [37]. Picard tools (v1.95) was then used to add read groups and mark duplicates [http://broadinstitute.github.io/picard/]. The mpileup program within Samtools carried out SNP calling (with options: -q20 -Q20 -AB −ugf) and these variants were then filtered using the bcftools package within Samtools (v0.1.18).

### Annotation of genome variants

Variants distinguishing the genomes of the F_6_ parental birds were classified as exonic - synonymous or non-synonymous, intronic, 5’-upstream, 3’-downstream or intergenic, as determined by the SNPEff program (v3.6c) [38]. Non-synonymous coding variants were further designated as predicted to be highly deleterious to protein function (henceforth, “potentially functional mutations”), moderately deleterious or having a low likelihood of being deleterious to protein function. In all, a total of 5,718,725 - 6,154,628 variants (in all 5 males v all 5 females) were annotated within each sample. Some of the identified variants were used to test the QTLR, as described below.

### 600K high density SNP genotyping of F_6_ birds

F_6_ birds were genotyped genome-wide with the high density 600k Affymetrix SNP array [32] for GWAS. DNA was available from 1,615 MDV challenged birds. After quality control (QC), 1,192 F_6_ birds provided high quality genotypes (178, 234, 357, 221 and 202 birds from Families 1 to 5, respectively). Genotyping was performed using 200 ng of gDNA with the standard protocol for the Axiom Affymetrix platform (Affymetrix). Samples were amplified using the Axiom 2.0 Reagent Kit (#901758, Thermo Fisher Scientific), and the resultant product checked for both quantity and quality of fragmentation. QC was performed using absorbance assay for quantity and running a sample on a 4% agarose E-Gel (#G800804, Thermo Fisher Scientific) to check for fragmentation. The samples were then hybridised to the Axiom Chicken Genotyping array (#902148, Thermo Fisher Scientific). Hybridisation, wash, stain and scanning were carried out within the GeneTitan MC Instrument. The resulting .CEL files were loaded into Axiom Analysis Suite (v2.0) for first stage analysis.

### GWAS of the F_6_ population to identify MD resistance QTL

The F_6_ genotypes were used to map QTL that impacted survival following MDV challenge. After applying Affymetrix standard QC the remaining markers were further filtered to remove markers with minor allele frequency ≤ 0.01 and markers with significant deviation from Hardy–Weinberg Equilibrium (P ≤ 0.001). An independent GWAS was then carried out within each of the 5 families of this study, using JMP Genomics SNP-Trait association Trend test (JMP Genomics, Version 9, SAS Institute Inc., Cary, NC, 1989-2019). Survival to the end of the experiment was taken as the Censor Variable; age at death was taken as a survival trait and MHC as a class variable and fixed effect. To obtain experimental significance thresholds, the Proportion of False Positives (PFP) [39,40] was used to correct for multiple tests.

### Identification of QTL and their confidence intervals

Although many marker x family combinations were nominally significant to highly significant (comparison wise P ≤ 0.05 to P ≤ 0.001), very few remained significant after PFP corrections for multiple tests. Nevertheless, visual inspection of the chromosomal Manhattan plots by families showed distinct clusters of markers with high −LogP values intermixed with markers with low −LogP values (**Figure 1**). Therefore, following Lipkin et al. [41], we identified QTL by using a moving average of −LogP (mAvg) to smooth the Manhattan plots. We used a window size of ~0.1 Mb (27 markers) with step size of 1 marker and a critical threshold mAvg of −LogP = 2.0 (P = 0.01) to declare significance, and Log drop 1 [42] to define QTLR boundaries.

**Figure 1:**
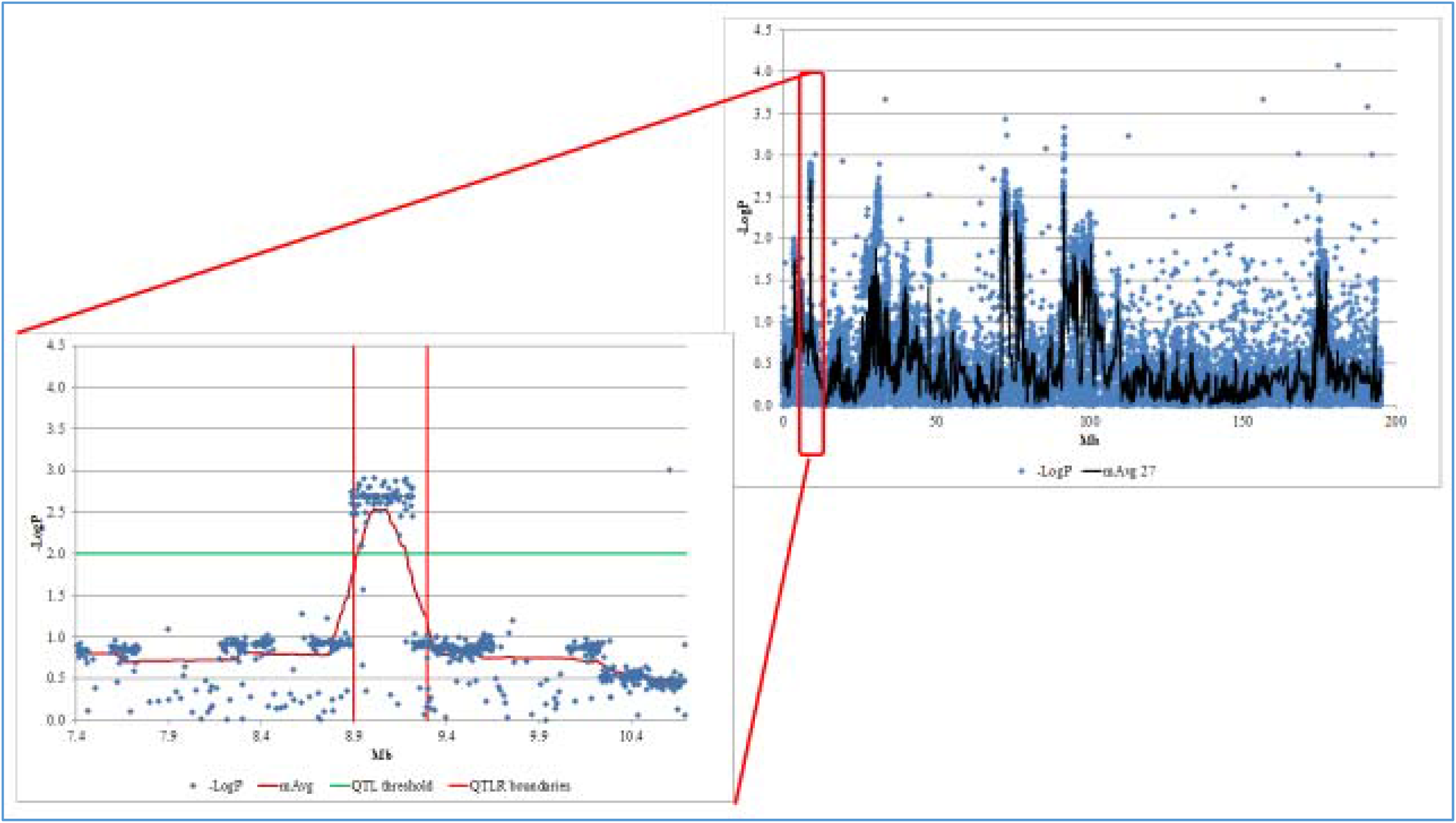
Manhattan plot of a significant F_6_ QTLR (Family 3: QTLR 1, Chromosome 1). LogP, −Log_10_P of a single marker; mAvg, moving average of −LogP values of a window of 27 markers; QTLR boundaries, boundaries of the QTL region obtained by Log Drop 1 (see Methods).

Many QTLR had overlapping boundaries across two or more families, thus providing replication and increased assurance of significance. Conservatively assuming such overlaps represents the same underlying genetic element, we combined these QTLR within and across families, taking the start and end log-drop boundaries of the QTLR as the first upstream and last downstream marker across all families in the QTLR. **Table 1** shows final list of 38 QTLR after consolidation within and across families. Also shown are the number of genes (Ensembl database v79) and non-coding RNAs [43] found in each QTLR

**Table 1.**
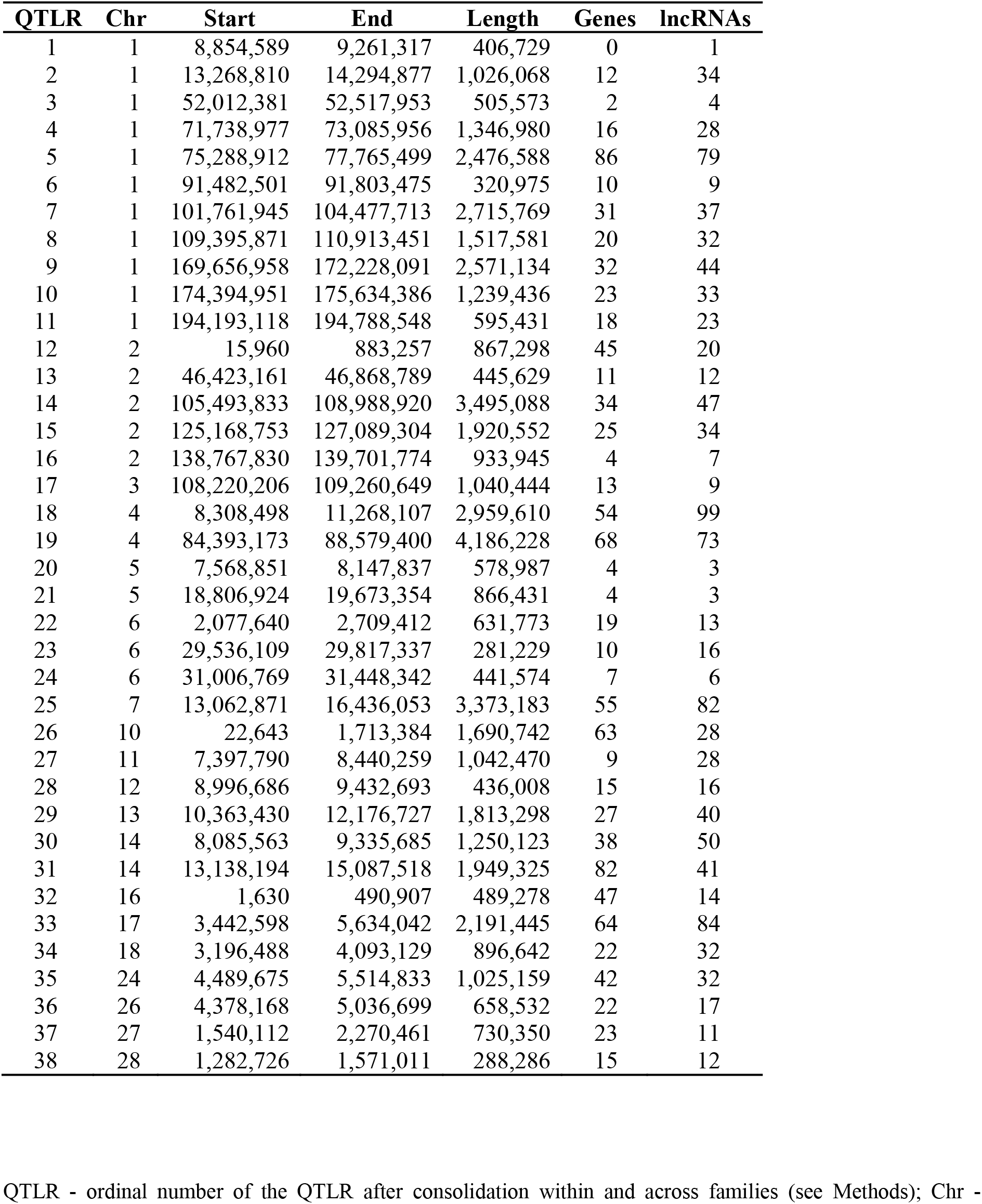
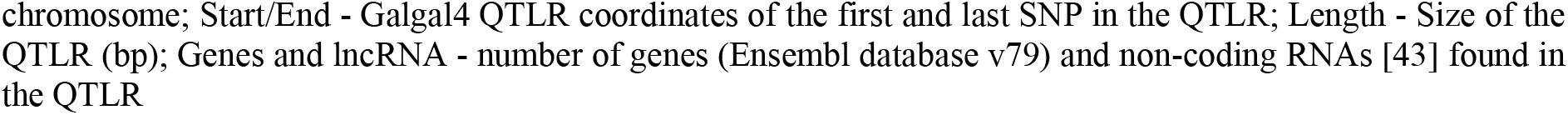
Mapped QTLRs.

| QTLR | Chr | Start | End | Length | Genes | lncRNAs |
| --- | --- | --- | --- | --- | --- | --- |
| 1 | 1 | 8,854,589 | 9,261,317 | 406,729 | 0 | 1 |
| 2 | 1 | 13,268,810 | 14,294,877 | 1,026,068 | 12 | 34 |
| 3 | 1 | 52,012,381 | 52,517,953 | 505,573 | 2 | 4 |
| 4 | 1 | 71,738,977 | 73,085,956 | 1,346,980 | 16 | 28 |
| 5 | 1 | 75,288,912 | 77,765,499 | 2,476,588 | 86 | 79 |
| 6 | 1 | 91,482,501 | 91,803,475 | 320,975 | 10 | 9 |
| 7 | 1 | 101,761,945 | 104,477,713 | 2,715,769 | 31 | 37 |
| 8 | 1 | 109,395,871 | 110,913,451 | 1,517,581 | 20 | 32 |
| 9 | 1 | 169,656,958 | 172,228,091 | 2,571,134 | 32 | 44 |
| 10 | 1 | 174,394,951 | 175,634,386 | 1,239,436 | 23 | 33 |
| 11 | 1 | 194,193,118 | 194,788,548 | 595,431 | 18 | 23 |
| 12 | 2 | 15,960 | 883,257 | 867,298 | 45 | 20 |
| 13 | 2 | 46,423,161 | 46,868,789 | 445,629 | 11 | 12 |
| 14 | 2 | 105,493,833 | 108,988,920 | 3,495,088 | 34 | 47 |
| 15 | 2 | 125,168,753 | 127,089,304 | 1,920,552 | 25 | 34 |
| 16 | 2 | 138,767,830 | 139,701,774 | 933,945 | 4 | 7 |
| 17 | 3 | 108,220,206 | 109,260,649 | 1,040,444 | 13 | 9 |
| 18 | 4 | 8,308,498 | 11,268,107 | 2,959,610 | 54 | 99 |
| 19 | 4 | 84,393,173 | 88,579,400 | 4,186,228 | 68 | 73 |
| 20 | 5 | 7,568,851 | 8,147,837 | 578,987 | 4 | 3 |
| 21 | 5 | 18,806,924 | 19,673,354 | 866,431 | 4 | 3 |
| 22 | 6 | 2,077,640 | 2,709,412 | 631,773 | 19 | 13 |
| 23 | 6 | 29,536,109 | 29,817,337 | 281,229 | 10 | 16 |
| 24 | 6 | 31,006,769 | 31,448,342 | 441,574 | 7 | 6 |
| 25 | 7 | 13,062,871 | 16,436,053 | 3,373,183 | 55 | 82 |
| 26 | 10 | 22,643 | 1,713,384 | 1,690,742 | 63 | 28 |
| 27 | 11 | 7,397,790 | 8,440,259 | 1,042,470 | 9 | 28 |
| 28 | 12 | 8,996,686 | 9,432,693 | 436,008 | 15 | 16 |
| 29 | 13 | 10,363,430 | 12,176,727 | 1,813,298 | 27 | 40 |
| 30 | 14 | 8,085,563 | 9,335,685 | 1,250,123 | 38 | 50 |
| 31 | 14 | 13,138,194 | 15,087,518 | 1,949,325 | 82 | 41 |
| 32 | 16 | 1,630 | 490,907 | 489,278 | 47 | 14 |
| 33 | 17 | 3,442,598 | 5,634,042 | 2,191,445 | 64 | 84 |
| 34 | 18 | 3,196,488 | 4,093,129 | 896,642 | 22 | 32 |
| 35 | 24 | 4,489,675 | 5,514,833 | 1,025,159 | 42 | 32 |
| 36 | 26 | 4,378,168 | 5,036,699 | 658,532 | 22 | 17 |
| 37 | 27 | 1,540,112 | 2,270,461 | 730,350 | 23 | 11 |
| 38 | 28 | 1,282,726 | 1,571,011 | 288,286 | 15 | 12 |

QTLR - ordinal number of the QTLR after consolidation within and across families (see Methods); Chr -
chromosome; Start/End - Galgal4 QTLR coordinates of the first and last SNP in the QTLR; Length - Size of the QTLR (bp); Genes and lncRNA - number of genes (Ensembl database v79) and non-coding RNAs [43] found in the QTLR

### RNA-Seq analysis

#### MDV challenge experiment

An experiment measuring host response to MDV viral strain 686 challenge (based on gene expression measured by a whole genome RNA-seq assay) was carried out with 10 male and 10 female birds from HL W36 commercial production hybrids. The parental lines used to produce the W36 commercial hybrid are closely related to the lines used to develop the F_6_ experimental cross. Previously it was reported that males are more resistant to MD than females [44–46]. On this basis, we compared Differentially Expressed Genes (DEGs) of males and females as surrogate for comparison of chicks from more- and less-resistant lines. Hence, gender balanced groups were used for the RNA-seq challenge experiment, with 5 males + 5 females in each of the challenged and control groups. This allowed us to identify viral response genes in each sex. Thus, this experiment included 4 comparisons: Females - challenged v control; Males - challenged v control; Controls - males v females; Challenged - males v females. Although some of the DEGs in the cross-sex comparisons will be due to sex rather than response to MDV, others can be expected to reveal some of the genes behind the host immune responses and potential differential susceptibility.

To reflect infection in the field, ten 6-day old HL W36 commercial birds (5 males + 5 females) were infected with 500 pfu of the very virulent plus (vv+) MDV strain 686, by subcutaneous injection in the neck (virus kindly provided by the USDA/Avian Disease and Oncology Lab, East Lansing, Michigan). All birds were vaccinated at 1 day of age with HVT/SB1 vaccine. Spleen tissues from these 10 challenged and from 10 unchallenged control birds were harvested at 4 dpi, flash frozen in liquid nitrogen and stored at −80°C for subsequent RNA preparation and RNA-Seq analysis. All birds were housed together, with challenged and control chicks separated by 2.44 meters.

#### RNA preparation

RNA was prepared from the 4-dpi flash-frozen spleen tissues of each of the above 20 chicks after homogenization in Qiazol reagent (Qiagen, Manchester, UK), and subsequent preparation using RNeasy RNA isolation kit (Qiagen, Manchester, UK) as per the manufacturer’s guidelines. RNA was resuspended in dH_2_O.

#### Transcriptomic sequencing and analysis

Strand-specific 100 bp paired-end RNA-Seq was carried out by GATC (Konstanz, Germany) using an Illumina HiSeq2500 genome sequencer. 19,510,400 – 26,596,800 reads representing at least 15x coverage were produced for each sample. Sequencing quality was assessed by means of FastQC [35] and reads mapped to the reference genome (Galgal4) with Tophat2 (v2.0.14) [47] using the Bowtie2 aligner (v2.2.3) [48]. Untrimmed reads with stated insert sizes were mapped using the Ensembl 79 genome annotation. 78.1% of reads mapped to the Galgal4 reference genome. The Cuffdiff package within Cufflinks (v2.2.1) was used to quantify transcripts and determine differential expression between experimental groups [49].

### QTLR tests

#### Selection of candidate genomic elements underlying the 38 QTLR for further testing and validation in the 8 HL elite pure lines

In principal a random selection of markers from a QTLR region could be used for testing in the 8 elite-sire validation populations. However, we decided to take a step forward, and search the QTLR for attractive candidate elements that could tested in the validation populations. The assumption being that for a marker to show association across a number of populations, it must be in high LD with the causative mutation in all of these populations. For this to hold, the marker must be very close to the causative mutation, which is what we would expect for markers taken from the quantitative trait element itself.

#### Identification of candidate genes within QTLR

The BioMart data mining tool within the Ensembl database (release 79) was used to identify genes lying under the 38 QTLR identified from the F_6_ data (**Table 1**). This information was then compared with the gene expression data (from RNA-Seq analysis of the MDV challenge experiment) in order to identify potential candidate genes for MD resistance. The DAVID analysis tool (v6.8) (https://david.ncifcrf.gov/home.jsp) identified gene ontology (GO) terms associated with DEGs under each QTLR. Following this, Ingenuity Pathway Analysis software (IPA v2.3) (https://www.qiagenbioinformatics.com/products/ingenuity-pathway-analysis/) revealed which canonical pathways were perturbed by MDV infection in the host, and allowed us to analyze the gene interaction networks involved in the host response. Likely upstream regulators of differentially regulated genes were identified and DEGs in male and female birds were compared.

#### Selection of genetic markers within QTLR for Genotyping

Markers located in genomic elements underlying each QTLR (genes, miRNAs, lncRNAs, and potentially functional mutations) were identified from genome sequences aligned to the reference chicken genome for each of the 8 elite lines utilized in this study. SNPs predicted to alter amino acids were preferentially chosen, but variants were also selected to encompass the entire gene where possible. All SNPs selected were validated using 200 samples from cohorts separated by 13-15 generations from each line to confirm that the marker was polymorphic (i.e. identified both homozygote and heterozygote individuals), and to define the minimum number of markers required to define a haplotype within each genomic element (gene, miRNA, or lncRNA) in each line. This minimum number of SNPs was then used to identify the haplotype status of each individual bird. The elite lines were selected for MD resistance, but the selection was at a low level, as selection emphasis was for production-related traits including egg numbers, shell quality and feed efficiency. If SNPs were homozygous in both generations, then the SNP composition was assumed monomorphic for the intervening and subsequent generations.

#### Selection of candidate coding genes

Twenty candidate coding genes (**Table 2**) were selected for further study in the 8 elite sire lines based on (i) position under a QTLR defined by this or previous genetic studies, (ii) known biology of the gene (e.g. involvement in innate immunity, cell death, or cancer), (iii) polymorphisms (presence of variants), (iv) differential expression between challenged and control, or more and less resistant birds - either in this or previous studies (e.g. Smith et al 2011), and (v) allele specific expression [50]. Specific details for the detection assays are provided in **S1 Table**.

**Table 2.**
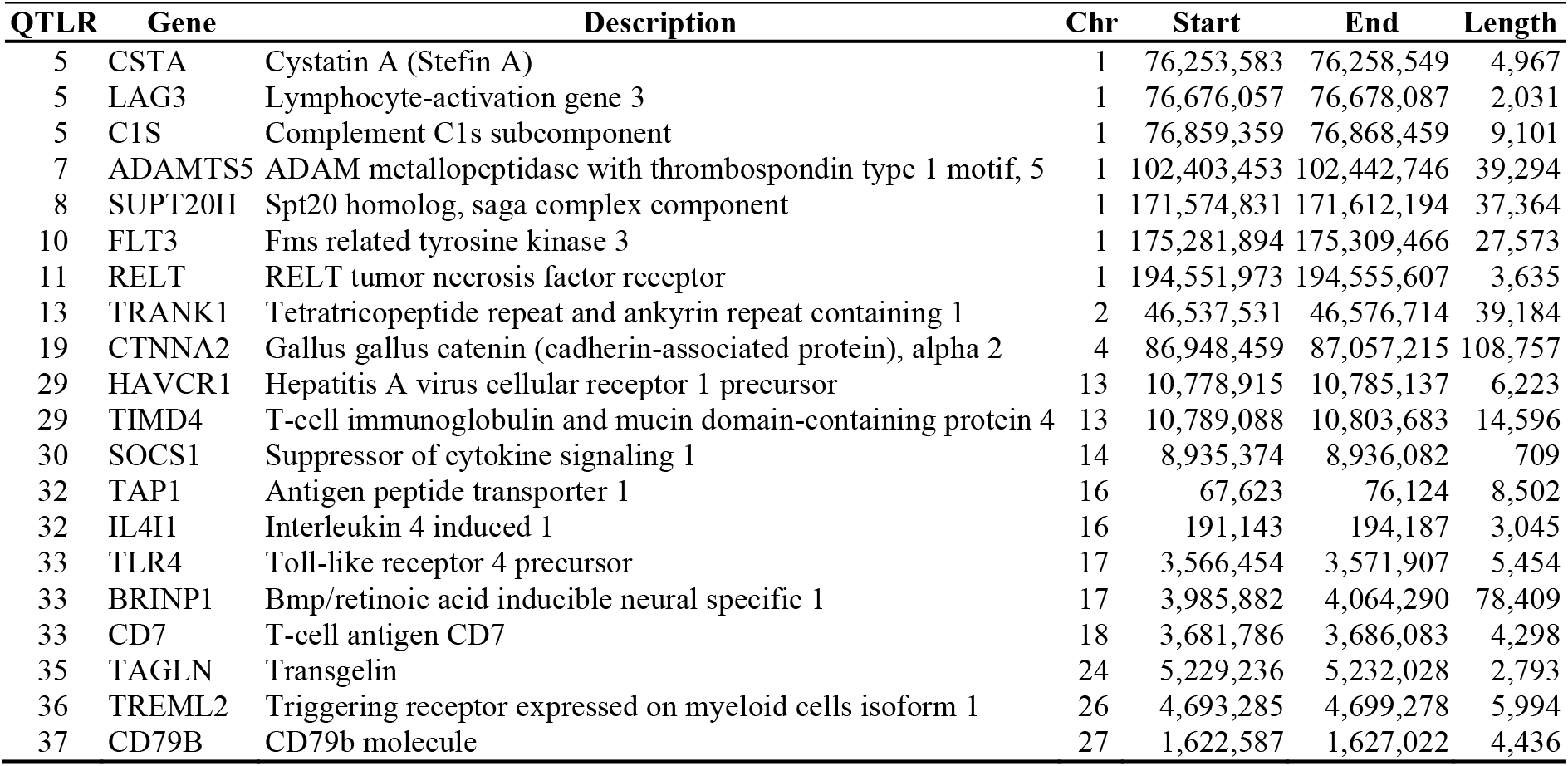
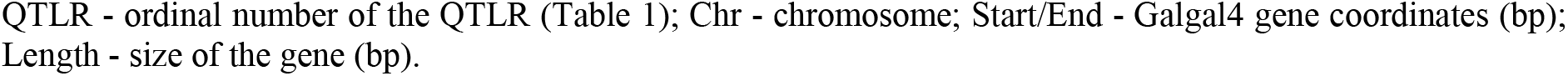
Candidate genes in the QTLRs, by location (Galgal4).

| QTLR | Gene | Description | Chr | Start | End | Length |
| --- | --- | --- | --- | --- | --- | --- |
| 5 | CSTA | Cystatin A (Stefin A) | 1 | 76,253,583 | 76,258,549 | 4,967 |
| 5 | LAG3 | Lymphocyte-activation gene 3 | 1 | 76,676,057 | 76,678,087 | 2,031 |
| 5 | C1S | Complement C1s subcomponent | 1 | 76,859,359 | 76,868,459 | 9,101 |
| 7 | ADAMTS5 | ADAM metalloproteinase with thrombospondin type 1 motif, 5 | 1 | 102,403,453 | 102,442,746 | 39,294 |
| 8 | SUPT20H | Spt20 homolog, saga complex component | 1 | 171,574,831 | 171,612,194 | 37,364 |
| 10 | FLT3 | Fms related tyrosine kinase 3 | 1 | 175,281,894 | 175,309,466 | 27,573 |
| 11 | RELT | RELT tumor necrosis factor receptor | 1 | 194,551,973 | 194,555,607 | 3,635 |
| 13 | TRANK1 | Tetratricopeptide repeat and ankyrin repeat containing 1 | 2 | 46,537,531 | 46,576,714 | 39,184 |
| 19 | CTNNA2 | Gallus gallus catenin (cadherin-associated protein), alpha 2 | 4 | 86,948,459 | 87,057,215 | 108,757 |
| 29 | HAVCR1 | Hepatitis A virus cellular receptor 1 precursor | 13 | 10,778,915 | 10,785,137 | 6,223 |
| 29 | TIMD4 | T-cell immunoglobulin and mucin domain-containing protein 4 | 13 | 10,789,088 | 10,803,683 | 14,596 |
| 30 | SOCS1 | Suppressor of cytokine signaling 1 | 14 | 8,935,374 | 8,936,082 | 709 |
| 32 | TAP1 | Antigen peptide transporter 1 | 16 | 67,623 | 76,124 | 8,502 |
| 32 | IL4I1 | Interleukin 4 induced 1 | 16 | 191,143 | 194,187 | 3,045 |
| 33 | TLR4 | Toll-like receptor 4 precursor | 17 | 3,566,454 | 3,571,907 | 5,454 |
| 33 | BRINP1 | Bmp/retinoic acid inducible neural specific 1 | 17 | 3,985,882 | 4,064,290 | 78,409 |
| 33 | CD7 | T-cell antigen CD7 | 18 | 3,681,786 | 3,686,083 | 4,298 |
| 35 | TAGLN | Transgelin | 24 | 5,229,236 | 5,232,028 | 2,793 |
| 36 | TREML2 | Triggering receptor expressed on myeloid cells isoform 1 | 26 | 4,693,285 | 4,699,278 | 5,994 |
| 37 | CD79B | CD79b molecule | 27 | 1,622,587 | 1,627,022 | 4,436 |

QTLR - ordinal number of the QTLR (Table 1); Chr - chromosome; Start/End - Galgal4 gene coordinates (bp); Length - size of the gene (bp).

#### Selection of candidate non-coding RNAs

With many causal variants for quantitative traits known to reside within regulatory elements [51], we deemed it important to examine non-coding transcripts as well. In contrast to the candidate genes, these were selected more or less on the basis of position underlying QTLR alone, with no information on potential biological function.

##### Micro RNAs (miRNAs)

A total of 46 miRNAs were found underlying the QTLR. Sequence information was used to identify markers segregating in HL lines. For some miRNAs, polymorphisms were not found, while for others multiple polymorphic sites were observed. Since most of the miRNAs are very short, haplotype information could be derived only for some of them. Ultimately, 17 miRNAs were chosen for further association analysis (**Table 3**). Detection assays for 30 SNPs lying within these 17 transcripts were developed (**S1 Table**).

**Table 3.** Candidate miRNAs in the QTLRs, by location (Galgal4).

| QTLR | Gene Name | Ensembl Gene ID | Chr | Start | End | Length |
| --- | --- | --- | --- | --- | --- | --- |
| 5 | gga-mir-6553 | ENSGALG00000026406 | 1 | 76,381,420 | 76,381,519 | 100 |
| 7 | gga-mir-1738 | ENSGALG00000025522 | 1 | 104,085,133 | 104,085,223 | 91 |
| 12 | gga-mir-1749 | ENSGALG00000025364 | 2 | 569,326 | 569,423 | 98 |
| 12 | gga-mir-6710 | ENSGALG00000027384 | 2 | 724,388 | 724,497 | 110 |
| 18 | gga-mir-1462 | ENSGALG00000025207 | 4 | 8,488,386 | 8,488,495 | 110 |
| 25 | gga-mir-1659 | ENSGALG00000025154 | 7 | 13,177,026 | 13,177,126 | 101 |
| 25 | gga-mir-6624 | ENSGALG00000026751 | 7 | 15,758,919 | 15,759,028 | 110 |
| 25 | gga-mir-1570 | ENSGALG00000025238 | 7 | 16,274,218 | 16,274,317 | 100 |
| 26 | gga-mir-6660 | ENSGALG00000028738 | 10 | 1,076,742 | 1,076,851 | 110 |
| 30 | gga-mir-1644 | ENSGALG00000025220 | 14 | 8,124,516 | 8,124,601 | 86 |
| 31 | gga-mir-6643 | ENSGALG00000027935 | 14 | 13,143,450 | 13,143,559 | 110 |
| 31 | gga-mir-1588 | ENSGALG00000025419 | 14 | 14,125,510 | 14,125,611 | 102 |
| 31 | gga-mir-1715 | ENSGALG00000025576 | 14 | 14,285,835 | 14,285,936 | 102 |
| 35 | gga-mir-6612 | ENSGALG00000026265 | 24 | 4,779,927 | 4,780,036 | 110 |
| 35 | gga-mir-6671 | ENSGALG00000027111 | 24 | 5,035,783 | 5,035,892 | 110 |
| 35 | gga-mir-1745 | ENSGALG00000026550 | 24 | 5,163,387 | 5,163,473 | 87 |
| 37 | gga-mir-6573 | ENSGALG00000028331 | 27 | 1,532,553 | 1,532,671 | 119 |

QTLR - ordinal number of the QTLR (Table 1); Chr - chromosome; Start/End - miRNA bp coordinates, Length - size of the miRNA (bp).

##### Long non-coding RNAs (lncRNAs)

With a view to testing some lncRNAs potentially associated with MDV resistance, five small QTLR were chosen which harboured only a few (1-6) lncRNAs. From these RNAs, up to 5 lncRNAs per QTLR were tested for association with MD. Thus, there was relatively high likelihood that one of these would be the genomic element housing the causative mutation. From among these, a further 17 transcripts were chosen for further association analysis (**Table 4**). Two lncRNAs overlapped (labelled with negative distance). Detection assays for 80 SNPs across these 17 transcripts were developed (**S1 Table**).

**Table 4.** Candidate lncRNA in the QTLRs, by location (Galgal4).

| QTLR | gene id; transcript id; stable transcript id | Chr | Start | End | Length | Distance | Strand | Exons |
| --- | --- | --- | --- | --- | --- | --- | --- | --- |
| 1 | R192;R192.1;RI_1_8950428_8950983_1_1 | 1 | 8,950,428 | 8,950,983 | 556 |  | + | 1 |
| 3 | R1291;R1291.1;RI_1_52122457_52122949_1_1 | 1 | 52,122,457 | 52,122,949 | 493 | 43,171,475 | - | 1 |
| 3 | R1292;R1292.1;RI_1_52183105_52322961_6_1 | 1 | 52,183,105 | 52,322,961 | 139,857 | 60,157 | - | 6 |
| 3 | R1293;R1293.2;RI_1_52481252_52482976_1_1 | 1 | 52,481,252 | 52,482,976 | 1,725 | 158,292 | - | 1 |
| 3 | R1293;R1293.3;RI_1_52481282_52488508_2_1 | 1 | 52,481,282 | 52,488,508 | 7,227 | -1,693 | - | 2 |
| 20 | R12773;R12773.1;RI_5_7604020_7604920_1_1 | 5 | 7,604,020 | 7,604,920 | 901 |  | + | 1 |
| 20 | R12779;R12779.1;RI_5_7727117_7728010_1_1 | 5 | 7,727,117 | 7,728,010 | 894 | 122,198 | - | 1 |
| 20 | R12784;R12784.1;RI_5_7960049_7961460_1_1 | 5 | 7,960,049 | 7,961,460 | 1,412 | 232,040 | + | 1 |
| 21 | R13147;R13147.6;RI_5_18814123_18815319_2_1 | 5 | 18,814,123 | 18,815,319 | 1,197 | 10,852,664 | + | 2 |
| 21 | R13148;R13148.1;RI_5_18861679_18862530_1_1 | 5 | 18,861,679 | 18,862,530 | 852 | 46,361 | + | 1 |
| 21 | R13149;R13149.1;RI_5_19632618_19793846_5_1 | 5 | 19,632,618 | 19,793,846 | 161,229 | 770,089 | - | 5 |
| 24 | R15316;R15316.1;RI_6_31138889_31157672_3_1 | 6 | 31,138,889 | 31,157,672 | 18,784 |  | - | 3 |
| 24 | R15317;R15317.1;RI_6_31199730_31200561_1_1 | 6 | 31,199,730 | 31,200,561 | 832 | 42,059 | - | 1 |
| 24 | R15318;R15318.1;RI_6_31204510_31210163_7_1 | 6 | 31,204,510 | 31,210,163 | 5,654 | 3,950 | + | 7 |
| 24 | R15319;R15319.1;RI_6_31263768_31264516_1_1 | 6 | 31,263,768 | 31,264,516 | 749 | 53,606 | + | 1 |
| 24 | R15320;R15320.2;RI_6_31277294_31279639_1_1 | 6 | 31,277,294 | 31,279,639 | 2,346 | 12,779 | - | 1 |
| 24 | R15321;R15321.1;RI_6_31368941_31369400_1_1 | 6 | 31,368,941 | 31,369,400 | 460 | 89,303 | - | 1 |

#### Selection of potentially functional variants in coding genes

Analysis of the complete sequences of the 10 F_0_ birds showed multiple SNPs distinguishing the founder birds that could impact gene function. Alternative variants fixed in one parental line compared to the other and which were predicted to cause highly deleterious changes in the gene protein product, were chosen as candidates for further study. Across the genome as a whole, a total of 252 variants of this nature were identified in 217 genes, with 22 of these variants lying within genes underlying the 38 QTLR. Twelve of these variants (representing 9 genes) were segregating in one or more of the 8 elite lines and were chosen for further studies in these lines. These 12 SNPs are summarized in **Table 5**. Specific details on the detection assays are provided in **S1 Table**.

**Table 5.**
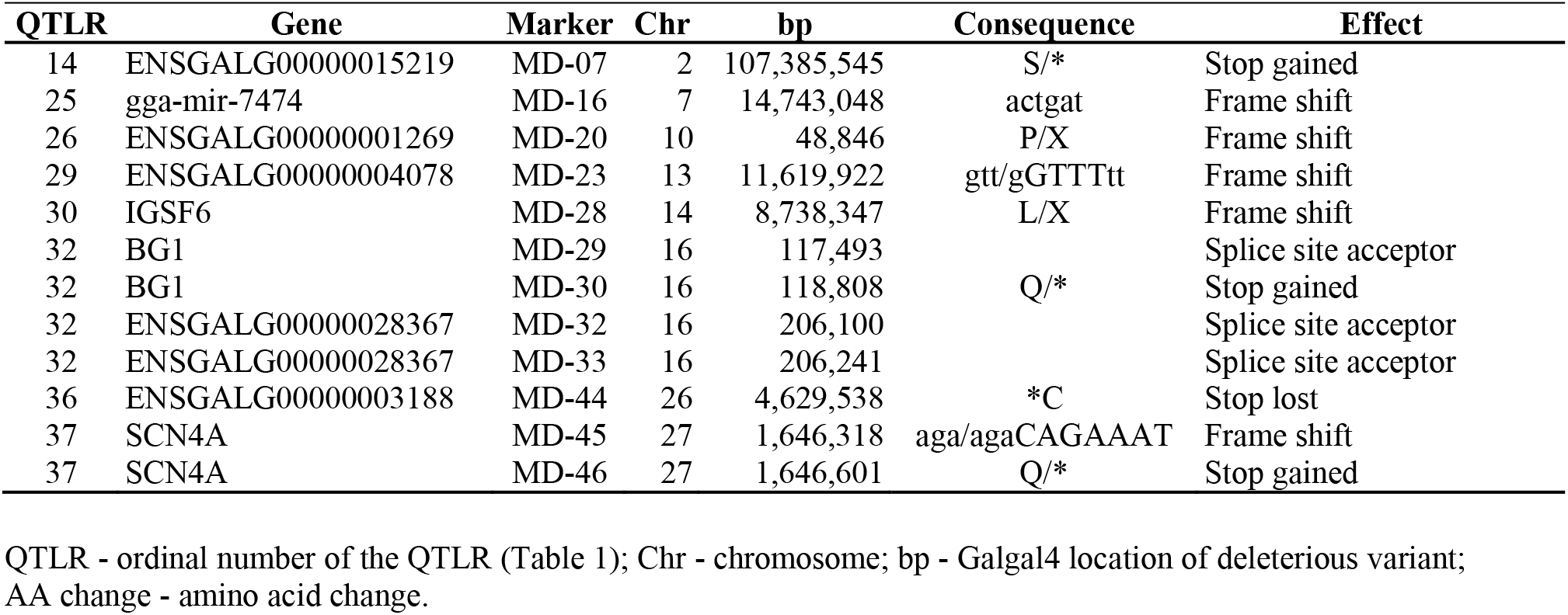
Markers in the QTLRs with genetic variations predicted to be functionally deleterious to the gene product.

| QTLR | Gene | Marker | Chr | bp | Consequence | Effect |
| --- | --- | --- | --- | --- | --- | --- |
| 14 | ENSGALG00000015219 | MD-07 | 2 | 107,385,545 | S/* | Stop gained |
| 25 | gga-mir-7474 | MD-16 | 7 | 14,743,048 | actgat | Frame shift |
| 26 | ENSGALG00000001269 | MD-20 | 10 | 48,846 | P/X | Frame shift |
| 29 | ENSGALG00000004078 | MD-23 | 13 | 11,619,922 | gtt/gGTTTtt | Frame shift |
| 30 | IGSF6 | MD-28 | 14 | 8,738,347 | L/X | Frame shift |
| 32 | BG1 | MD-29 | 16 | 117,493 |  | Splice site acceptor |
| 32 | BG1 | MD-30 | 16 | 118,808 | Q/* | Stop gained |
| 32 | ENSGALG00000028367 | MD-32 | 16 | 206,100 |  | Splice site acceptor |
| 32 | ENSGALG00000028367 | MD-33 | 16 | 206,241 |  | Splice site acceptor |
| 36 | ENSGALG00000003188 | MD-44 | 26 | 4,629,538 | *C | Stop lost |
| 37 | SCN4A | MD-45 | 27 | 1,646,318 | aga/agaCAGAAAT | Frame shift |
| 37 | SCN4A | MD-46 | 27 | 1,646,601 | Q/* | Stop gained |

QTLR - ordinal number of the QTLR (Table 1); Chr - chromosome; bp - Galgal4 location of deleterious variant; AA change - amino acid change.

#### Selecting QTLR markers from the candidate genomic elements, for genotyping in the 8 elite HL pure lines

Markers located in the genomic elements underlying the QTLR (genes, miRNAs, lncRNAs, and singleton variants with potentially functional effects), were identified from genome sequences aligned to the reference chicken genome for each of the lines utilized in this study. For the candidate genes underlying the QTLR, SNPs predicted to alter amino acids were preferentially chosen. For all genomic elements (not including the singleton functional mutations), variants were also selected to encompass the entire gene when possible, so as to define haplotype status of each bird for subsequent association tests. All SNPs selected were validated using the same approach as defined above

### Genotyping of the eight elite lines

Genomic DNA was extracted from whole blood using salt/ethanol precipitation and stored at −20°C until use. DNA was diluted to 25 ng/ul. Genotyping was done as single-plex assays on the SNPline system (LGC, UK). All assays used KASP chemistry, which is a fluorescent-based competitive allele-specific detection method [52]. Genotyping was done in 1,536 well plates, and results analysed with Kraken software (LGC, Hoddesdon, UK) as previously described [53]. Specific details on primers for each assay are provided in **S1 Table**.

### QTLR association analysis

Association of markers and marker-haplotypes with sires progeny-tested for post-challenge survival in the 8 HL elite lines was tested using JMP Genomics SNP-Trait association Trend test (JMP Genomics, Version 9, SAS Institute Inc., Cary, NC, 1989-201). When analysing a line-marker combination, the generation and MHC background were taken as class variables and fixed effects, and when analysing a marker combining all lines together, the line was added to the fixed effects. To obtain experimental significance thresholds, the Proportion of False Positives (PFP) [39,40] was used to correct for multiple tests.

## Results

### High resolution MD QTL mapping in Advanced Full-Sib inter-cross F_6_ families

Following genotyping of the F_6_ families by the Affymetrix 600K HD SNP chip, 40-43% of SNPs (depending on family) were found to be informative and passed QC (total, 238,777 to 259,098 informative SNPs per family; average of 246,400 SNPs in a family). The distribution of the marker P-values was fairly uniform over the range of allele frequencies. A clear excess of small P-values (an indication of presence of true QTL effects) was not seen. In accordance with this, only in families 2, 4 and 5, were markers significant at PFP ≤ 0.2. However, n_1_ - the estimated number of falsified null hypotheses at PFP ≤ 0.2 [39,40], ranged from 1,718 in Family 1 (0.7% of all markers tested in the family) to 17,385 in Family 2 (6.7% of markers tested), indicating the actual presence of true QTL effects.

As described in Methods, many markers were nominally significant, and distinct clusters of nominally significant markers intermixed with non-significant markers were clearly seen. Hence, to identify QTL we used a moving average of marker −Log_10_P values (mAvg) across a window size of 0.1 Mb (27 SNPs) to smooth the plots. Examining Manhattan plots (**Figure 1** and **S1 Figure**), showed that the smoothing did yield a good monotonic plot, allowing identification of QTL and the use of the drop method [42] to define their boundaries.

Using a threshold of mAvg ≥ 2.0 and QTLR boundaries defined by Log Drop 1, a total of 57 regions meeting our criteria for significance were identified, ranging from 4 to 16 in a family. In some families, several regions clustered within 1 Mb of one another. Nine such chromosome-by-family clusters were identified, containing 2-4 significant sub-regions per cluster. Conservatively, we counted each cluster as representing a single within-family QTLR. Similarly, aligning QTLR maps of the five families, showed that 6 significant regions overlapped or were within 1 Mb of one another across two families. Here too, counting the same QTLR across different families only once, resulted in a final total of 38 QTLR being identified (**Table 1**). These 38 QTLR were the focus of further examination of variants as defined by multiple methodologies employed below.

### Annotation of QTLR

Transcripts of genes located under the 38 mapped QTLR were identified by using the Biomart tool within the Ensembl database (v79) (**S2 Table**). The number of known annotated genes in each QTLR ranged from 0 to 86, for a total of 1,072 genes. Using the annotation developed and described by [53], which was based on characterisation of full-length transcripts using long-read sequencing and a bioinformatics pipeline to define coding and non-coding RNAs, we identified 1 to 99 lncRNAs within each defined QTLR, with a total 1,153 lncRNAs (**Table 1**).

### Identification and Analysis of SNPs in F_0_ parental birds

To investigate variants residing in regions of the genome potentially associated with MDV resistance, the 10 founder parents that gave rise to the F_6_ population used in this study were sequenced and variants determined between males and females. Combined across both lines 8,273,112 SNPs were identified. Examining the 38 QTLR within the genome highlighted 133,394 SNPs in these regions, which were fixed for alternative SNP variants with one parental line compared to the other (**S3 Table**). With a view to testing for causality, any of these variants could then be tested for genetic association with the progeny-test for survival in the 8 elite HL lines.

### Transcriptomic analysis of male and female commercial birds challenged with MDV

Based on the observation noted above, that male and female birds tend to show differential resistance, we examined the expression profiles of five male and five female W36 birds challenged with vv^+^MDV, and 5 male and 5 female W36 non-challenged control birds (see Methods). Gene expression results from this transcriptomic analysis are presented in **S4 Table**.

First, we compared gene expression in challenged and control chicks within each sex of W36 birds; 185 genes (58 up- and 127 down-regulated) were differentially expressed in male chicks, while there were 114 genes (62 up- and 52 down-regulated) differentially expressed upon challenge of the female birds. When these genes are compared across sex, the individual responses are seen to be quite different, with minimal overlap of DEGs between the male and female groups (**Figure 2**).

**Figure 2:**
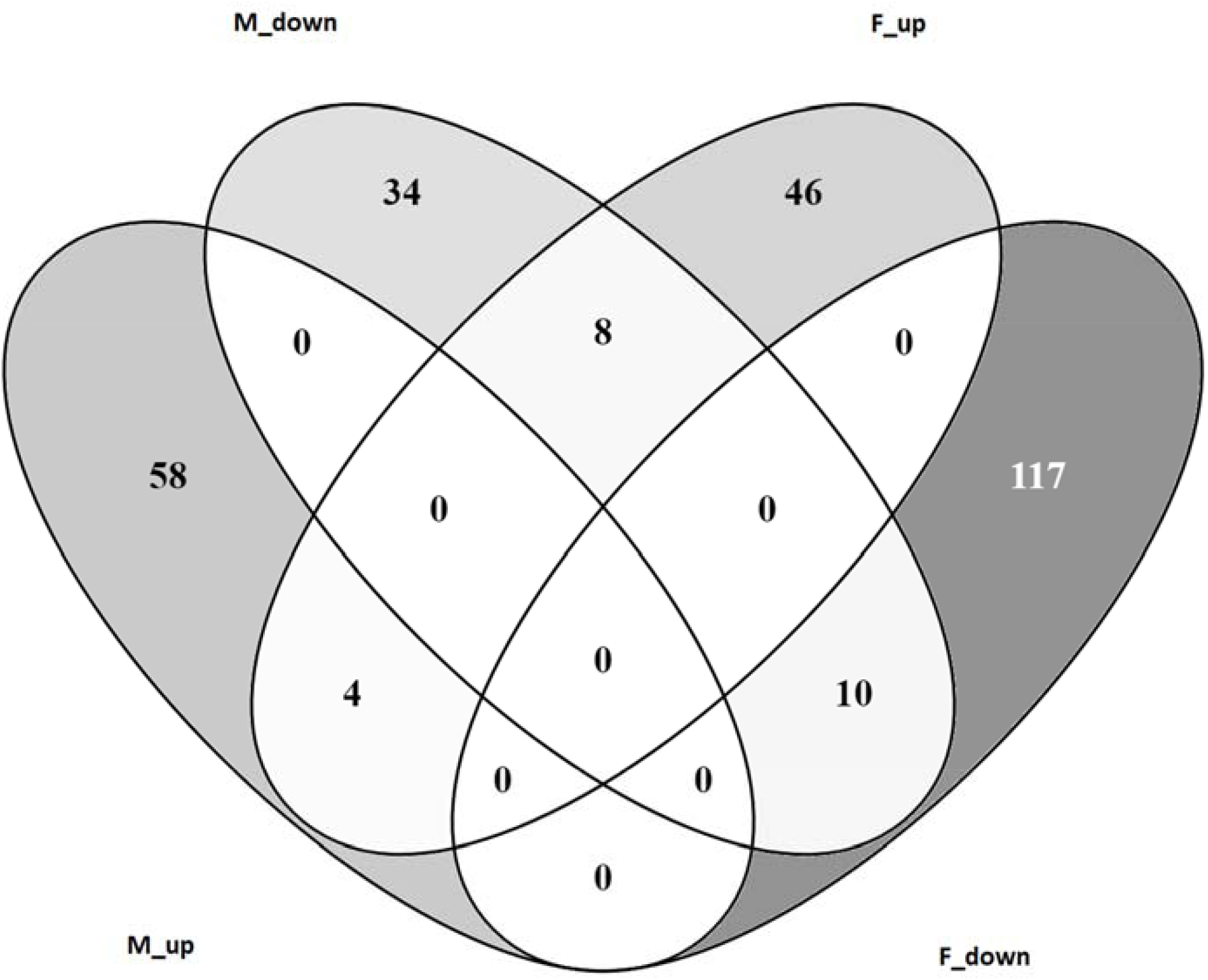
Venn diagram showing the intersection of host responses in male and female W36 birds. M-up – number of genes up regulated in male birds; M-down - number of genes down regulated in male birds; F-up – number of genes up regulated in female birds; F-down - number of genes down regulated in female birds.

When the different responses were compared, it is seen that biological pathways predominantly involved in the host response in female chicks include activation of cancer signalling, whereas the DEGs identified in the male birds have roles in the Th1 immune response and dendritic cell maturation amongst others (**Figure 3A**). The diseases and biofunctions associated with the genes involved in each response are depicted in **Figure 3B**. Strikingly, with MD being an oncogenic disease, functions related to cancer and tumorigenesis are seen to be up-regulated in female birds, but are repressed in the male chicks. Investigation of genes acting as upstream regulators of identified DEGs again indicates differential regulatory mechanisms in each sex. *TGFB* appears to be a key regulator of mechanisms in females, whereas *STAT1* (an interferon response gene) is suggested to have a more prominent role in the males (**Figure 3C**).

**Figure 3:**
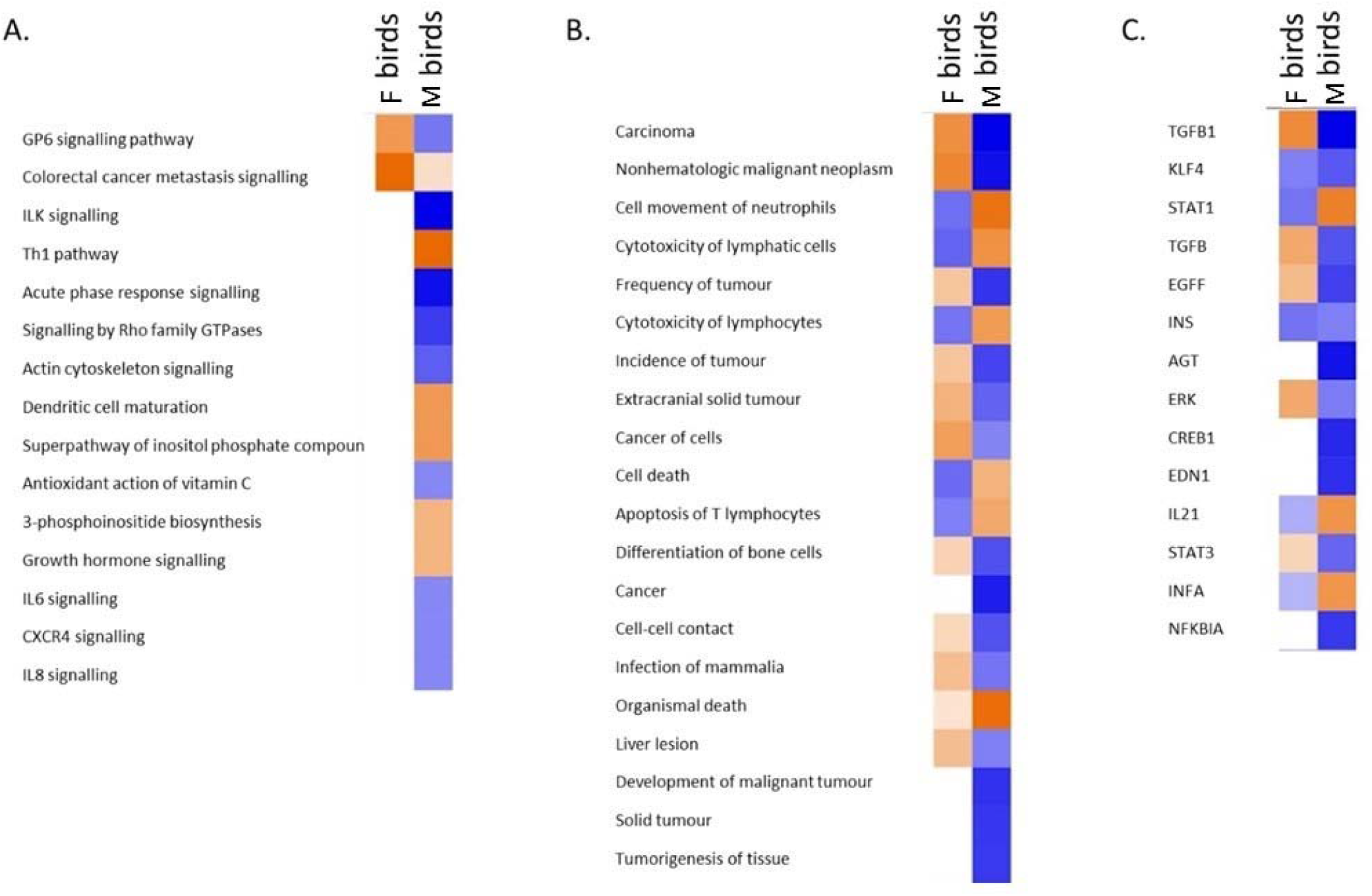
Comparison of the gene expression underlying the host response in W36 birds. Orange colours represent up-regulation while blue indicates down-regulation. Differential activation/repression is indicated as shown: (**A**). Biological pathways (**B**). Diseases and biofunctions (**C**). Upstream regulators.

Examination of the unchallenged control birds from each group (male and female W36 birds) identified 302 genes which were expressed in an inherently different manner between the sexes. So, these DEGs, being independent of MD challenge, are due to gender differences and/or intrinsic immune differences, manifesting in differing resistance to infection. Significant DEGs include the obvious immune related genes but also various serine proteases and carboxypeptidases, which may suggest a role in resistance to MDV. Genes previously implicated in MDV resistance including MHC genes, *IRG1* [18] and *SCYC1* [16], were also highlighted.

### Analysis of differentially expressed genes located in QTLR

When DEGs are located within QTLR affecting MD resistance, it is plausible that these genes may be the causative quantitative trait gene (QTG) responsible for the QTLR effect. Based on this hypothesis, the list of genes located in the 38 QTLR identified in this study was compared to the list of DEGs. Genes that were both expressed under challenge and that were also located under one of the 38 mapped QTL, were considered likely candidate QTG (**S5 Table**).

To determine if any particular types of genes were represented by the DEGs underlying significant genomic regions (identified either from this or previous studies [31,33,54–56]), the gene ontologies associated with these DEGs were studied. **S6 Table** shows that immunoglobulins, stimulus response genes and genes involved in regulating the immune response are all highly represented. Analysis of the biological pathways controlled by these genes was also examined. To look for possible common pathways, network analysis was performed. Three networks stood out significantly: cell movement, cell signalling and cancer (**Figure 4A**); cell signalling and nervous system development (**Figure 4B**) and antimicrobial resistance, inflammation and cell death (**Figure 4C**). When candidate upstream regulators of these genes were investigated, the cytokines *TNF*, *EDN1*, *IL1B* and *IFNG* were all indicated as potentially regulating many of the genes under study. Interestingly, the *TNF* gene is located in QTLR 18 on Chr 4, 387,071 bp downstream of the top window in this QTLR.

**Figure 4:**
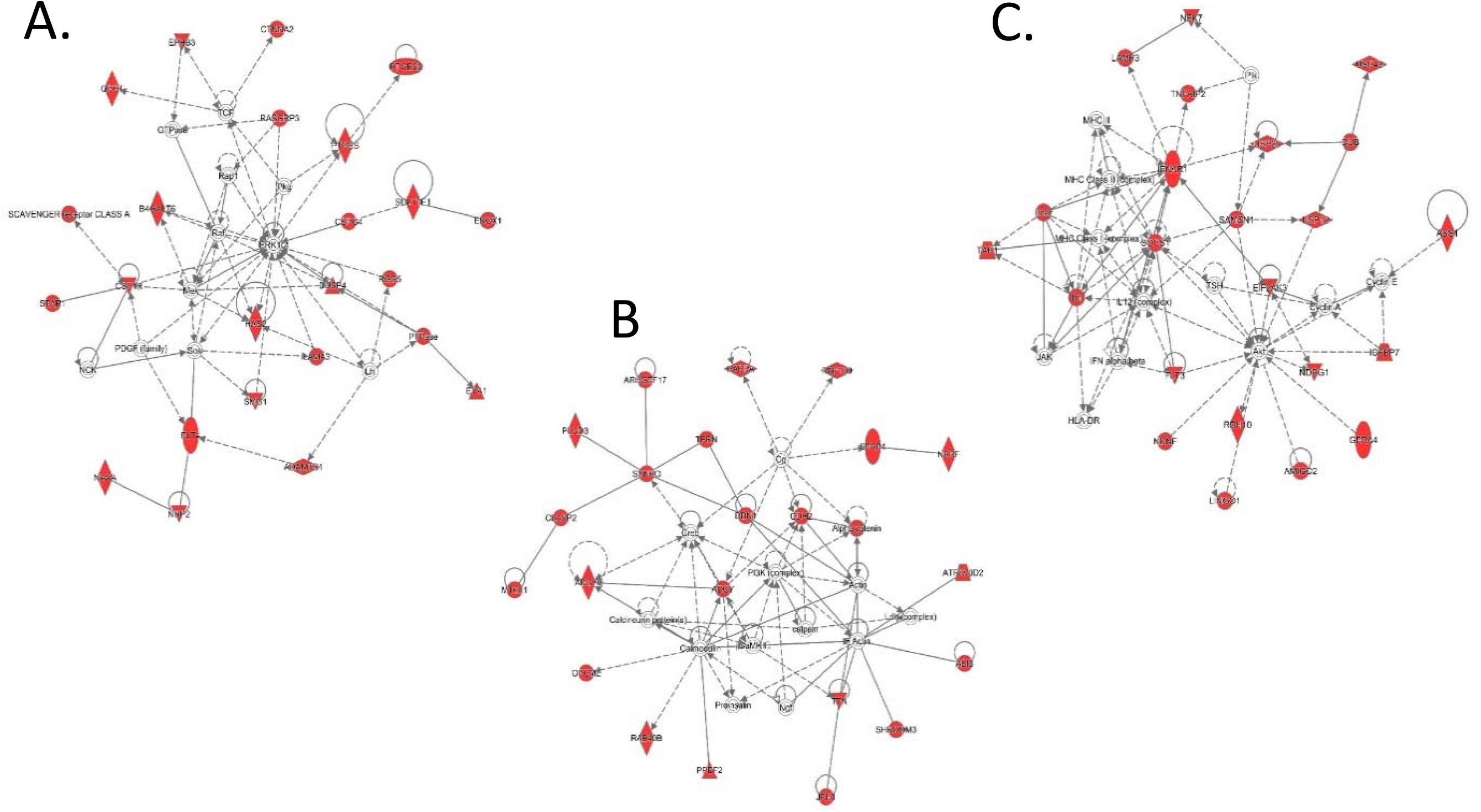
Gene networks represented by differentially expressed genes found under the 38 significant QTLR. Genes coloured in red are genes within the network which are in the dataset under study. (**A**). cell movement, cell signalling and cancer (**B**). Cell signalling and nervous system development and (**C**). Antimicrobial resistance, inflammation, and cell death.

### Validating QTLR for association with MDV resistance

#### Populations, genomic elements and markers

In a unique aspect of this study, QTLR identified in the F_6_ mapping population were re-tested for association with MD resistance in 8 pure lines maintained under selection for multiple commercial production traits at HL. Markers for QTLR validation were chosen from among the various classes of genomic elements identified underlying the QTLR, namely genes, miRNAs and lncRNAs. Also included were a group of singleton coding SNPs predicted to have deleterious effects on protein structure and function (the functional mutations).

#### Genetic association tests

When possible, a number of markers were genotyped in each candidate gene, so that association could be based on marker haplotypes rather than individual markers. When this is done, all of the information is in the ‘Haplotypes’. The individual markers are usually in complete LD and hence do not add anything to the analysis. Thus, for statistical analyses what is important are the number of element x line tests, rather than number of marker x line tests. We counted an element x line test as significant if the haplotype was significant, or if one or more markers in the element was significant.

A total of 355 markers located in 66 genetic elements (genes, miRNA, lncRNA, and functional mutations) underlying 26 of the 38 QTLR were tested for association with MD mortality in the 8 elite lines (**S7 Table)**. In 46 of the elements, multi-marker haplotypes were also tested. The association tests were carried out separately within each of the 8 elite lines, and also in data combined across the 8 lines, yielding 2,032 P-values. A total of 387 element by line tests were performed.

##### Significance of individual candidate genes

Twenty genes located in the QTLR were tested by a total of 127 gene x line tests of markers encompassing the entire gene, A test was considered significant if the haplotype x line test was significant, or if any one of the markers encompassing the haplotype were significant. All but *IL4I1*, *TAGLN* and *CD79B* (in QTLR 32, 35 and 37), were significant in two or more marker x line tests. Thus, 17 genes may be taken as strong candidates to be a QTG (**S8 Table**).

##### Association of markers located in miRNA

A total of 17 miRNAs were chosen (**Table 3**), and 28 markers tested for association (**S7 Table**) for a total of 96 element x line tests. Only 5 significant to highly significant P-values were obtained. (**S8 Table)**.

##### Significance of individual miRNAs

Significant results were found in only 5 out of the 17 miRNA tested. Most interesting was gg-mir-6553 with the highest proportion of significant tests among the miRNAs (**S8 Table)**. This miRNA is located in QTLR 5, along with 3 significant genes (*CSTA, LAG3, and C1S*). The results indicate two putative causative quantitative elements in this QTLR: one upstream in the region of *CSTA*, the other downstream in the region of *LAG3* and *C1S*.

##### Associations of markers located in lncRNAs

A total 17 lncRNAs were chosen (**Table 4**), and 80 markers were tested for association (**S7 Table)** in a total of 114 element x line tests, of which 9 were significant (**S8 Table**).

##### Significance of individual lncRNAs

Significant results were found with 9 of the 17 lncRNAs tested (**S8 Table**). QTLR worth mentioning are 20 and 21 on Chr5, both tested by lncRNAs only, are detailed in **S1 File**.

##### Associations of QTLR functional mutations in coding genes

Twelve potentially functional mutations were tested in a total of 50 element x line tests, of which 6 were significant (**Table 5**).

##### Significance of genes harbouring potentially functional mutations

Significant results were found in 5 of the 9 QTLR genes harbouring potentially functional mutations. In Line WPR1, the 2 markers in the gene *BG1* (markers MD-29 and MD-30 in QTLR 32 on Chr 16) had practically the same P values as the markers of the candidate gene *TAP1* (**S8 Table**), thus strengthening the results of that gene. It should be noted that *BG1* and *TAP1* lie within the chicken MHC, as noted above, a locus known to impact MDV resistance.

#### Summary of element x line tests

**S9 Table** summarises the element x line tests by type of element. Tests of the coding genes are outstanding in the high proportion of significant element x line tests; more than double that of the other genomic elements. Since the candidate genes were chosen on the basis of much more information than the other classes of elements, the high proportion of significant tests can be taken to support the proposition that the process by which the candidate genes were selected was effective and that an appreciable fraction of the chosen candidate genes are the actual Quantitative Trait Genes. This must be qualified however, as the distribution of the different classes of genomic elements among the QTLRs was not random, and thus may be biasing the results.

### Validation of QTLR

A total of 26 QTLR were tested for significance of element x line tests. Of these, eight did not have any significant element x line tests, 5 had one significant test and 14 had 2 to 6 significant element x line tests. Thus, 18 QTLR can be considered as validated. Of the unconfirmed QTLR, QTLRs 1, 3 and 24 were tested by lncRNAs only, QTLRs 12 and 18 by miRNAs only, QTLR 14 by functional mutations only, and QTLR 25 by both miRNA and functional mutations. More markers need to be tested in these QTLR to decide if the lack of significance is a result of lack of informativity (Type 2 error), or if indeed these QTLR are false positives (Type I error). More detail on the association found within each individual QTLR is presented in **S1 File.**

## DISCUSSION

QTL mapping in an FSIL F_6_ population phenotyped for survival in the face of MDV challenge, identified 38 QTLR distributed over 19 autosomes. Use of such a resource allowed for the identification of QTLR at a higher resolution than have been mapped previously, thus allowing for easier identification of potential candidate genes. The mapped QTLR, along with genomic sequences of the F_0_ founder individuals, and transcriptomic information from challenged and control birds, has allowed us to identify genes, miRNAs, lncRNAs and potentially functional mutations located under these QTLR as candidates for association with progeny mortality from Marek’s Disease. Genetic association studies in multiple elite lines have confirmed the significant effects of most of these candidates on MD. Here we will discuss the potential role of some of the most significant candidates.

Many of the genes we have associated with MD response in this study have biological roles clearly relevant to the pathogenesis of MDV infection. One of the primary targets of the virus are B-cells and genes known to be associated with B-cells, include two of our candidates - *CD7*, the T-cell leukaemia antigen which is involved in T-cell/B-cell interactions and the Toll-like receptor, *TLR4* which is found on the surface of B-lymphocytes.

After initial infection and a period of latency, T-cells become infected. Genes related to T-cell signalling pathways include our MD-associated *ADAMTS5*, *CD7*, *HAVCR1*, *LAG3*, *RELT*, *TIMD4* and *TREML2*. *ADAMTS5* (ADAM Metallopeptidase With Thrombospondin Type 1 Motif 5) encodes a metalloproteinase that plays an important role in inflammation and cell migration. It also has a critical role in T-lymphocyte migration from draining lymph nodes following viral infection. HAVCR1 - Hepatitis A Virus Cellular Receptor 1 (T-Cell Immunoglobulin Mucin Receptor 1) is a receptor for TIMD4. HAVCR1 plays a critical role in regulating immune cell activity, particularly regarding the host response to viral infection, while TIMD4 is a T-cell immunoglobulin involved in regulating T-cell proliferation and lymphotoxin signalling. *LAG3* (Lymphocyte-Activation Gene 3) belongs to the immunoglobulin superfamily and acts as an inhibitory receptor on activated T-cells. It negatively regulates the activation, proliferation and effector function of both CD8^+^ and CD4^+^ T-cells as well as mediating immune tolerance. RELT is a member of the TNF-receptor superfamily. It can activate the NF-kappaB pathway and selectively bind TNF receptor-associated factor 1 (TRAF1). This receptor acts via CD3 signalling to stimulate T-cell proliferation, suggesting its regulatory role in the immune response. TREML2 (Triggering Receptor Expressed On Myeloid Cells Like 2) is a cell surface receptor that may play a role in both the innate and adaptive immune responses. It acts as a counter-receptor for CD276, with interaction with CD276 on T-cells enhancing T-cell activation.

Once infection of T-cells has occurred, the disease can then proceed to become oncogenic. Once again, we see that many of our MD associated genes have functions that have been implicated in cancer, including *BG1* which encodes an Ig-superfamily type I transmembrane receptor-like protein that contains an immuno-receptor tyrosine-based inhibition motif (ITIM). BG1 has previously been documented as conferring MHC-associated resistance to MDV-induced lymphoma [57]. Other candidates include *BRINP1* (silenced in some bladder cancers), *CD7* (associated with leukaemia), *CSTA* (encodes a stefin that functions as a cysteine protease inhibitor, suggested as a prognostic tool for cancer), *FLT3* (mutations in this gene are common in acute myeloid leukaemia), and *SUPT20H* (a known tumour rejection antigen).

One of the main pathologies of MD is its effect on the nervous system, and so it is interesting to see that some of our MD associated genes are involved in the function/growth of neurons (*SCN4A* and *CTNNA2*). *SCN4A* (Sodium Voltage-Gated Channel Alpha Subunit 4) encodes one member of the sodium channel alpha subunit gene family involved in generation and propagation of action potentials in neurons and muscle. CTNNA2 (Catenin Alpha 2) is thought to be involved in regulation of cell-cell adhesion and differentiation in the nervous system. It is required for proper regulation of cortical neuronal migration and neurite growth.

The remaining MD associated genes are seen to have general roles as immune system genes: *C1S*, *TAP1* and *SOCS1*. *C1S* (Complement component 1S) encodes a serine protease component of the complement system which enhances the host antibody immune response. TAP1 (Transporter 1, ATP Binding Cassette Subfamily B Member) is involved in the transport of antigens from the cytoplasm to the endoplasmic reticulum for association with MHC class I molecules. *SOCS1* (Suppressor of Cytokine Signalling 1) encodes a protein which functions downstream of cytokine receptors, and takes part in a negative feedback loop to attenuate cytokine signalling. All of these candidate genes had more than one test significant at P ≤ 0.05.

Examination of these genes and their significance of association with MDV resistance across the elite lines indicates a few top candidates, namely: the cluster of genes in QTLR5 (*CSTA*, *C1S* and *LAG3*), *FLT3* in QTLR10, *CTNNA2* in QTLR19 and *TAP1* in QTLR32.

Genes identified in this analysis include many novel candidates for resistance as well as highlighting genes proposed in previous studies. For example *CD8B* (T-cell glycoprotein), *CTLA4* (immunoglobulin) and *CD72* (B-cell associated) are postulated as important lncRNA target genes by You et al. [24] and are found under our QTLRs (*CD8B* - QTLR19), and differentially expressed in our transcriptomic work (*CTLA4* and *CD72*). Similarly, *ATF2* (involved in carcinogenesis is found in QTLR25) was proposed as an important target for the miRNA gga-mir15b during MDV infection [25]. Also in QTLR25 we find gga-mir-10b, previously seen to be upregulated in the spleen during MDV infection [27]. Other potentially interesting miRNA targets include *PBEF1* (pre-B-cell enhancing factor) and *FCHSD2* (involved in endocytosis) [26] that lie under QTLR2 and 11, respectively. Further genes previously linked with MDV resistance include *GH1* (growth hormone) and *CD79B* (B-cell antigen), both of which lie under our QTLR37.

One of the significant aspects of this research is that it utilized large, commercial production-relevant lines, and the challenge virus is a very virulent ++ strain, frequently encountered by production birds in the field. In contrast, most previously published MDV resistance research utilizes specialized research lines, many of which are inbred, and selected for differential response to MDV. These studies utilized laboratory strains of the virus, for which commercial production birds now appear to be resistant. Furthermore, this study investigated MD resistance genes in three distinct breeds of chickens, White Leghorn, White Plymouth Rock and Rhode Island Red, not just one laboratory line. Moreover, these MD association studies replicated the results from the FSIL study increasing our confidence in the causal nature of the QTLR, and possibly the genes and variants in MDV resistance. The response to the virus was measured as mortality in a large progeny group (approximately 30 daughters) for over 9,000 sires, using pre-existing information that had been developed within a commercially relevant production trait breeding program. This unique approach increases the relevance of the results to application into a commercial breeding program, while simultaneously provides information on the underlying mechanism of general viral resistance applicable to not only birds, but also other species. This information can provide insights into mechanisms for improving resistance or lead to the development of improved commercial vaccines.

## CONCLUSIONS

Utilizing an FSIL F_6_ population of birds phenotyped for response to Marek’s Disease Virus infection has allowed us to map QTLR for disease resistance at high-resolution. Combining this with expression data from challenged and control birds, we have identified candidate genes, miRNAs, lncRNAs and potentially functional mutations which have been validated in genetic association tests with MD mortality in diverse, elite lines of poultry. This most comprehensive genetic study to date supplies us with variants in candidate genes that can now go on to be functionally tested for their utility in marker assisted selection, improved vaccine development and potential future gene editing strategies.

## LIST OF ABBREVIATIONS

DEG: differentially expressed gene
FSIL: full-sib inter-cross line
GWAS: genome wide association study
HAPS: haplotypes
HL: Hy-Line
HVT: herpesvirus of turkeys
mAvg: moving average of −LogP of a marker association test
MD: Marek’s Disease
MDV: Marek’s Disease Virus
MHC: major histocompatibility complex
PCR: polymerase chain reaction
PFU: plaque forming units
QC: quality control
QT: quantitative trait
QTG: quantitative trait gene
QTL: quantitative trait locus
QTLR: QTL region
SB1: non-oncogenic MD virus
vv+: very virulent plus
SNP: single nucleotide polymorphism

## ETHICS APPROVAL

All procedures carried out on the birds involved in this study were conducted in compliance with Hy-Line International Institutional Animal Care and Use Committee guidelines.

## AVAILABILITY OF DATA

Data have been submitted to the European Nucleotide Archive (ENA) at EMBL-EBI under study accession numbers PRJEB39142 (WGS) and PRJEB39361 (RNAseq).

## AUTHOR CONTRIBUTIONS

JS carried out whole genome sequence, variant and RNAseq analyses, downstream biological analyses and wrote the manuscript; EL and MS performed all association analyses; JF provided all experimental animals, DNA samples and individual genotypes; DB conceived and managed the project. All authors contributed to the interpretation of the results and edited and approved the final manuscript.

## ACKNOWLEDGEMENTS

The authors would like to thank Choon-Kiat Khoo, Richard Kuo and Lel Eory (Roslin) for invaluable bioinformatics assistance. Also requiring thanks are Ashlee R. Lund, Amy M. McCarron, Kara Pinegar-Maxon and Grant Liebe (Hy-line) for their excellent technical assistance with genotyping. We are also extremely grateful to Edinburgh Genomics (Edinburgh, UK) for carrying out the 600K SNP genotyping and the whole genome sequencing of the parental birds. We would also like to thank the GATC sequencing facility (Konstanz, Germany) for generating the transcriptomic data used in this study.

## Supplementary Files

S1 Table – Primers

S1 Figure – Manhattan plots of significant QTLR

S2 Table – Ensembl genes under QTLR

S3 Table – Variants under QTLR

S4 Table – Differentially expressed genes

S5 Table – QTLR genes also differentially expressed

S6 Table – Gene ontology terms

S7 Table – All markers used in association tests

S8 Table – Association test results (p-values) S9 Table – Summary of element x line tests

S1 File – Details of association by individual QTLR

